# Identifying, Protecting and Managing Stopover Habitats for Wild Whooping Cranes on U.S. Army Corps of Engineers Lakes

**DOI:** 10.1101/2020.12.30.424870

**Authors:** Chester McConnell

**Affiliations:** Friends of the Wild Whoopers 8803 Pine Run Spanish Fort, Alabama, 36527, USA

**Keywords:** Aransas-Wood Buffalo population, Great Plains, *Grus americana*, lake, migration, pond, stopover habitat, reservoir, USACE, wetland, whooping crane

## Abstract

The Whooping Crane *(Grus americana)* is one of North America’s most endangered species. There is only one wild, self-sustaining migratory population of Whooping Cranes, the Aransas–Wood Buffalo population (AWBP). The birds of the AWBP migrate 4,000 km twice each year between their nesting grounds in northern Canada and their wintering grounds on the Texas Gulf Coast. During migration, AWBP Whooping Cranes must land at suitable ponds or wetlands to forage, rest or roost. The Whooping Crane Recovery Plan, developed by federal wildlife agencies in Canada and the USA, calls for the protection and management of Whooping Crane stopover locations within the migration corridor. Although major stopover areas have been protected, many other smaller sites remain to be identified. However, the Recovery Plan offers no specific entity to identify, protect and manage the latter. To address these deficiencies in information and activity, Friends of the Wild Whoopers partnered with the United States Army Corps of Engineers (USACE) within the AWBP migration corridor to share information about Whooping Cranes and their habitat needs and identify potential stopover locations on USACE properties that could be protected and managed for cranes. This partnership identified 624 potential stopover sites on 34 USACE lakes, principally in North and South Dakota, Nebraska, Kansas, Oklahoma and Texas, with commitments to manage the habitats as resources allow.

## INTRODUCTION

The Whooping Crane (*Grus americana*) is one of North America’s most threatened species (reviewed in French et al., 2019). It is considered *endangered* in both Canada and the USA, and is similarly categorized as *endangered* on the International Union for Conservation of Nature’s Red List of Threatened Species. It is a large bird, North America’s tallest, and also an ‘umbrella species,’ which means that by preserving the Whooping Crane and its habitat, many other species of birds and non-avian wildlife will also benefit. Several decades of captive breeding and reintroduction efforts have not yet produced self-sustaining offshoot populations of Whooping Cranes in the United States.

There remains only one wild, self-sustaining migratory population of Whooping Cranes, the Aransas-Wood Buffalo population (AWBP). This population nests and raises its chicks in Canada’s Wood Buffalo National Park in northern Alberta and the Northwest Territories (April - October) and winters on or near Aransas National Wildlife Refuge in Texas (October - April). The birds of the AWBP migrate 4,000 km twice each year between their nesting and wintering areas (Kuyt, 1992). The migration route takes them through two prairie provinces (Alberta and Saskatchewan) and six principal states in the Great Plains (North Dakota, South Dakota, Nebraska, Kansas, Oklahoma and Texas) (**Figure 1**). During migration, Whooping Cranes must land at any suitable wetland area when they get tired, when severe weather occurs or before nightfall. These stopover sites are important because they provide cranes with foraging opportunities and safe nocturnal roosts. Pearse et al. (2017) used GPS data from tagged AWBP Whooping Cranes to categorize the stopover habitats in the Great Plains portion of the migration corridor as follows: 50% emergent wetlands (e.g., small ponds with herbaceous vegetation), 25% lacustrine wetlands (e.g., lakes, reservoirs, impoundments), 20% riverine, and 5%dryland (“sites without discernible surface water”, but rarely used for more than one night). Clearly, AWBP Whooping Cranes are highly dependent on wetland habitats during their twice-yearly long-distance migrations.

**Figure 1.**
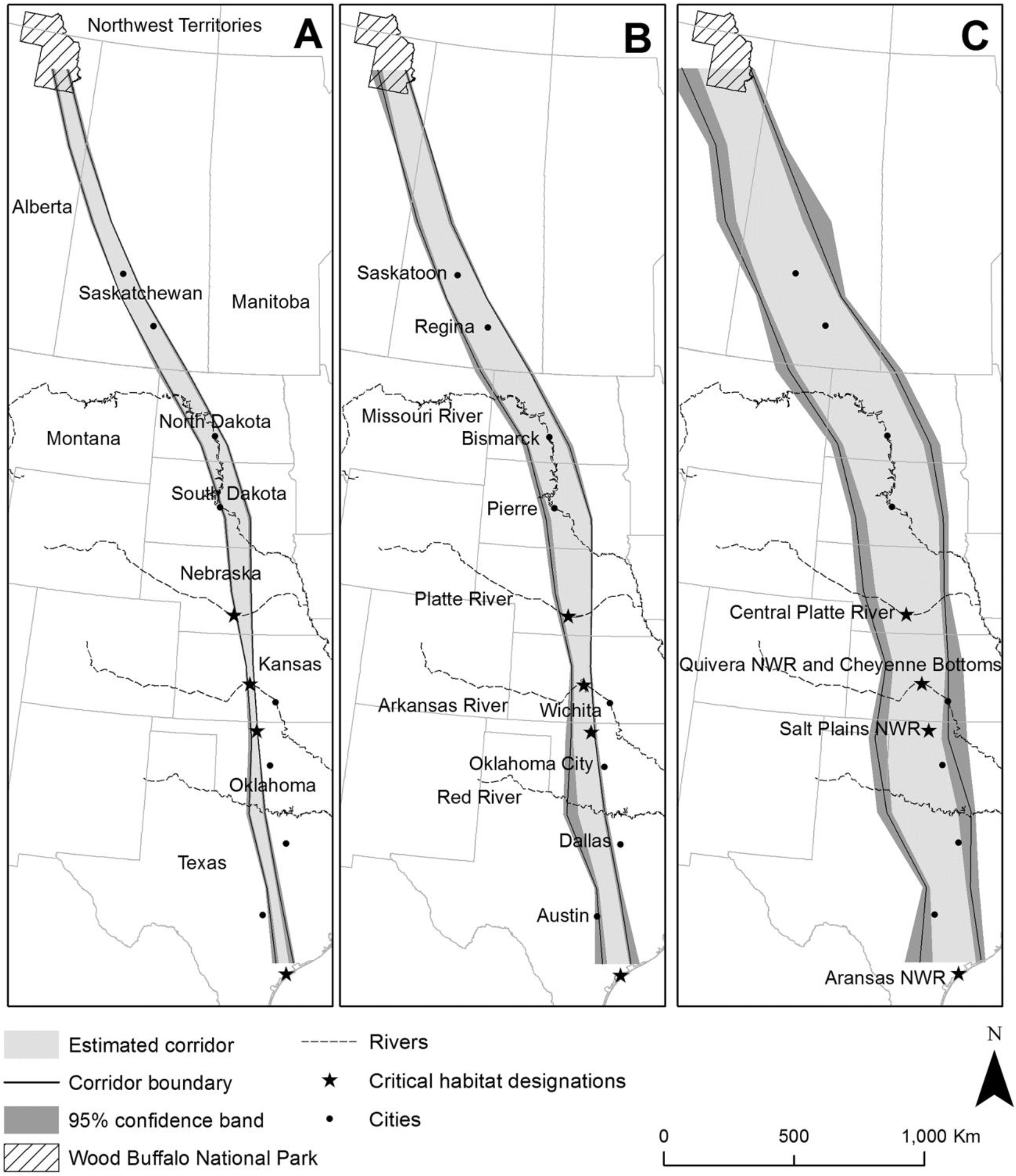
Migration corridors of Whooping Cranes of the Aransas-Wood Buffalo Population, showing the 50% core (A), 75% core (B), and 95% core migration areas, with 95% confidence bands [reproduced from Pearse et al. (2018), https://doi.org/10.1371/journal.pone.0192737, under the Creative Commons Universal Public Domain Dedication]. The illustrated corridors, running from the nesting area in Canada’s Wood Buffalo National Park to the wintering area at Aransas National Wildlife Refuge in Texas, are based on 75 years of compiled opportunistic sightings and 7 years of more recent GPS data of tagged Whooping Cranes (Pearse et al., 2018). Also indicated are areas designated as Whooping Crane *critical habitat* in the United States, and some cities and major rivers.

Since 1941, the AWBP has increased from 15 birds (Allen, 1952) to an estimated 506 as of winter 2019-2020 (United States Fish and Wildlife Service, 2020). Despite the increasing population size, the Whooping Cranes of the AWBP remain vulnerable to habitat destruction and gunshot. During the 200-year period from 1780 to 1980, wetland acreage in the Whooping Crane migration corridor within the United States declined by over 6 million ha (see Dahl, 1990, 2000). These habitats continue to be lost or degraded due to a variety of human activities, including wetland drainage (Samson et al., 2004), intensified farming and other changes to agricultural programs (Matson et al., 1997; Stehn and Pieto, 2008), and construction of wind energy facilities and power transmission lines (Pearse et al., 2016; Derby et al., 2018). Climate change is also likely to further reduce the stopover habitats available for Whooping Cranes (Chavez-Ramirez and Wehtje, 2012).

The Whooping Crane Recovery Plan (Canadian Wildlife Service and U.S. Fish and Wildlife Service, 2007) includes numerous references to wetlands known to be used as migration stopover sites. Important stopover sites in the United States include the Platte River bottoms near Kearney, Nebraska; Cheyenne Bottoms State Waterfowl Management Area and Quivira NWR in central Kansas; and Salt Plains NWR in northern Oklahoma (Figure 1C). These large sites have been designated as *critical habitat* for conservation of the Whooping Crane (United States Department of the Interior, 2017), but other stopover areas have also been identified, both large (Austin and Richert, 2001) and small (e.g., Pearse et al., 2017). Moreover, Whooping Cranes are not site-specific each migration and rarely use the same wetlands year to year (Pearse et al., 2018; 2020). Indeed, their selection of stopover locations may in part be influenced by year-to-year changes in wetlands availability (e.g., dependent on precipitation). Furthermore, there is evidence that Whooping Crane flock sizes may be increasing at some stopover locations, outpacing the overall growth of the AWBP, which may be an indicator of limited stopover habitat availability in those areas (Caven et al., 2020). Large aggregations of Whooping Cranes may increase the risk of catastrophic loss, e.g., from disease or adverse weather events (Caven et al., 2020). For these reasons, Friends of the Wild Whoopers (FOTWW), a 501(c)(3) organization, emphasizes that numerous other smaller stopover sites are also essential to ensure diverse opportunities for potential stopover use along the migration corridor.

As we noted previously (McConnell, 2018), the Whooping Crane Recovery Plan calls for the protection of existing wetlands as Whooping Crane stopover areas and the enhancement of those wetlands that have been degraded by woody plant encroachment, silting, and/or draining within the migratory corridor. An outline of recovery actions to achieve objectives is explained in the Recovery Plan (Canadian Wildlife Service and U.S. Fish and Wildlife Service, 2007). These actions include identifying, protecting, managing, and creating habitat. More specifically, the Recovery Plan (section 1.5.3.2.) highlights the need to “Ensure long-term protection of migration stopover sites. Work with landowners to ensure migration habitat remains suitable for cranes. Pursue stewardship agreements and conservation easements when needed, focusing on providing wetland mosaics” (page 49). However, the Recovery Plan offered no specific entity to protect and manage potential stopover sites.

Within the United States’ portion of the migratory corridor, FOTWW could find no ongoing concerted effort that focuses on protection or enhancement of many potential stopover areas (McConnell, 2018). Private conservation groups (e.g., Ducks Unlimited) and government agencies have played a significant role in protecting wetlands used by waterfowl and many other wildlife species throughout the AWBP Whooping Crane migration corridor. For example, funds from the sale of Duck Stamps have helped protect over 2.4 million ha of wetlands in the United States (National Wildlife Refuge Association, 2017), but many of those areas are managed for waterfowl in ways that may not be suitable for cranes (e.g., presence of tall emergent vegetation around the wetland perimeter or deeper water that would deter cranes from roosting). The most expensive part of establishing or improving habitat is land cost. If stopover habitat projects can be undertaken on government or tribal land (Indian Reservations), the cost would be relatively minimal. To address these deficiencies in information and activity, FOTWW initiated a survey of entities with large land holdings that could possibly provide additional stopover areas for migrating AWBP Whooping Cranes.

The first two phases of the project evaluated potential stopover habitat on 14 U.S. military bases and 7 Indian Reservations within the U.S. portion of the AWBP Whooping Crane migration corridor (McConnell, 2018). Here we report the results of phase 3, where FOTWW partnered with the U.S. Army Corps of Engineers (USACE) to evaluate Whooping Crane potential stopover habitats on USACE lake properties within the migration corridor (USACE districts Omaha, Kansas City, Tulsa, Fort Worth and Galveston). The USACE provides national leadership in the development, management, conservation and restoration of the nation’s water resources and provides real estate services for the agencies of the U.S. Department of Defense.

## METHODS

FOTWW and USACE developed a Memorandum of Understanding (MOU), effective 15 April 2018, to evaluate USACE lake properties for potential Whooping Crane stopover habitat. The project involves properties in six states through which the core-intensity Whooping Crane migration corridor passes — North Dakota, South Dakota, Nebraska, Kansas, Oklahoma and Texas — and one state, Montana, where low-intensity use by Whooping Cranes has been recorded (Figure 1) (Pearse et al., 2015; 2018; 2020). USACE lakes within the seven-state core migration corridor — there are 36 USACE lakes in total — are likely to become even more important to Whooping Cranes in the near future owing to the lakes’ prime locations and the managed water impoundment that ensures availability of wetlands habitat. These reservoirs will be especially vital when other stopover sites are lost to drought caused by climate change.

Included as part of the MOU (an unclassified USACE document) were the following conservation goals:

### Article IV - Understanding of the parties

- The USACE and the FOTWW desire to conserve freshwater, estuarine and coastal water resources, and natural communities inhabited by Whooping Cranes and other associated native wildlife. (Section 1)
- The USACE and the FOTWW desire to promote innovative thinking about conservation needs of Whooping Cranes to maintain healthy water resources and associated natural communities. (Section 2)
- Subject to the availability of resources and in accordance with applicable laws, regulations, Army policies, and FOTWW policies; the USACE and the FOTWW desire to conduct habitat assessments, develop recommendations, and conduct demonstration projects to improve Whooping Crane stopover habitat, roosting habitat and wintering habitat. (Section 5)

### Article V - Responsibilities

- The USACE and the FOTWW will cooperate in identifying opportunities to promote the conservation and/or restoration of Whooping Crane stopover habitat, water resources and natural ecosystems both on a project-specific level and on a national level along the migration corridor of the Whooping Cranes, consistent with the USACE mission and authorities to protect water resources. These opportunities may include identifying possible stopover habitat, surveying during the migration season for the presence of Whooping Cranes, developing Whooping Crane stopover habitat and other efforts to assist the USACE in executing its responsibilities under its authorities. (Section 5) The criteria used by FOTWW to identify suitable Whooping Crane stopover habitat were as per McConnell (2018), as follows:
- Lake, pond, wetland at least 0.12 ha;
- Lake, pond, wetland with a shallow area 12-25 cm deep for roosting;
- Glide path (for Whooping Cranes to land near the water body) is clear of obstructions (e.g., power lines);
- No thick vegetation or trees near the landing site: open landscapes allow Whooping Cranes to easily locate the ponds and provide for ready observation of any predator threats;
- Gradual or gentle slope into the water where it is shallow;
- Little or no emergent/submerged vegetation in the potential roost area;
- Extensive horizontal visibility from the potential roost site;
- At least 275 m from human development or disturbance.

Prior to visiting each USACE lake property, FOTWW analyzed satellite images (Google Earth) to identify locations of potential stopover habitat for Whooping Cranes, by applying the above criteria. Numerous candidate stopover locations were identified in this way for subsequent evaluation on the ground. The field trips allowed FOTWW not only to engage with local ‘lake managers’ and biologists about Whooping Crane biology and conservation needs, but also to ground truth the locations we had viewed on the satellite imagery. On-site interviews with lake personnel as well as FOTWW observations made during the lake evaluations informed our understanding of any ongoing wildlife habitat management programs. Some land and water management reports were also provided to FOTWW. Site visits were conducted by vehicle or by boat (n=8; see Table 1) during daylight and typically lasted 8-10 hours.

**Table 1.**
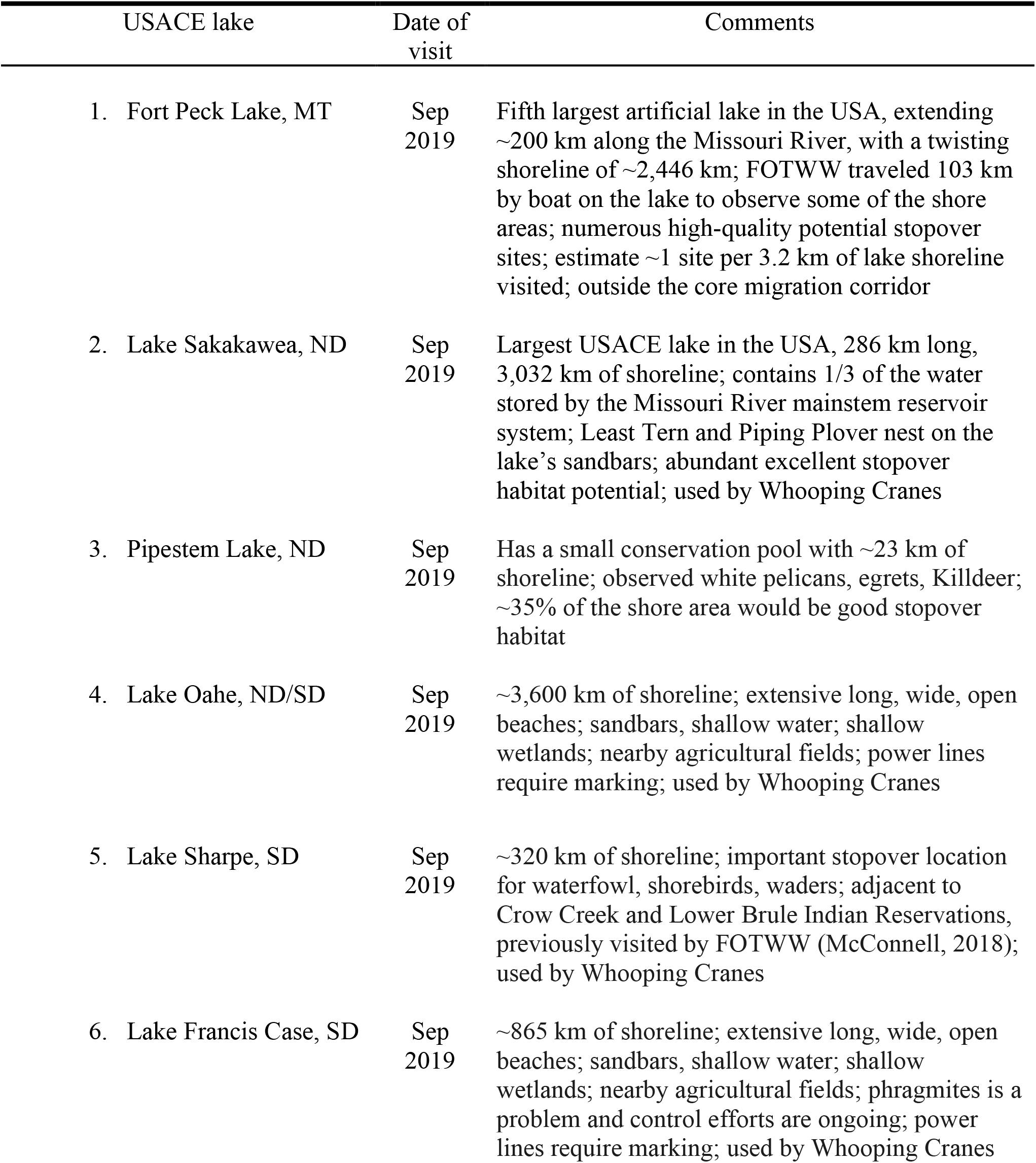
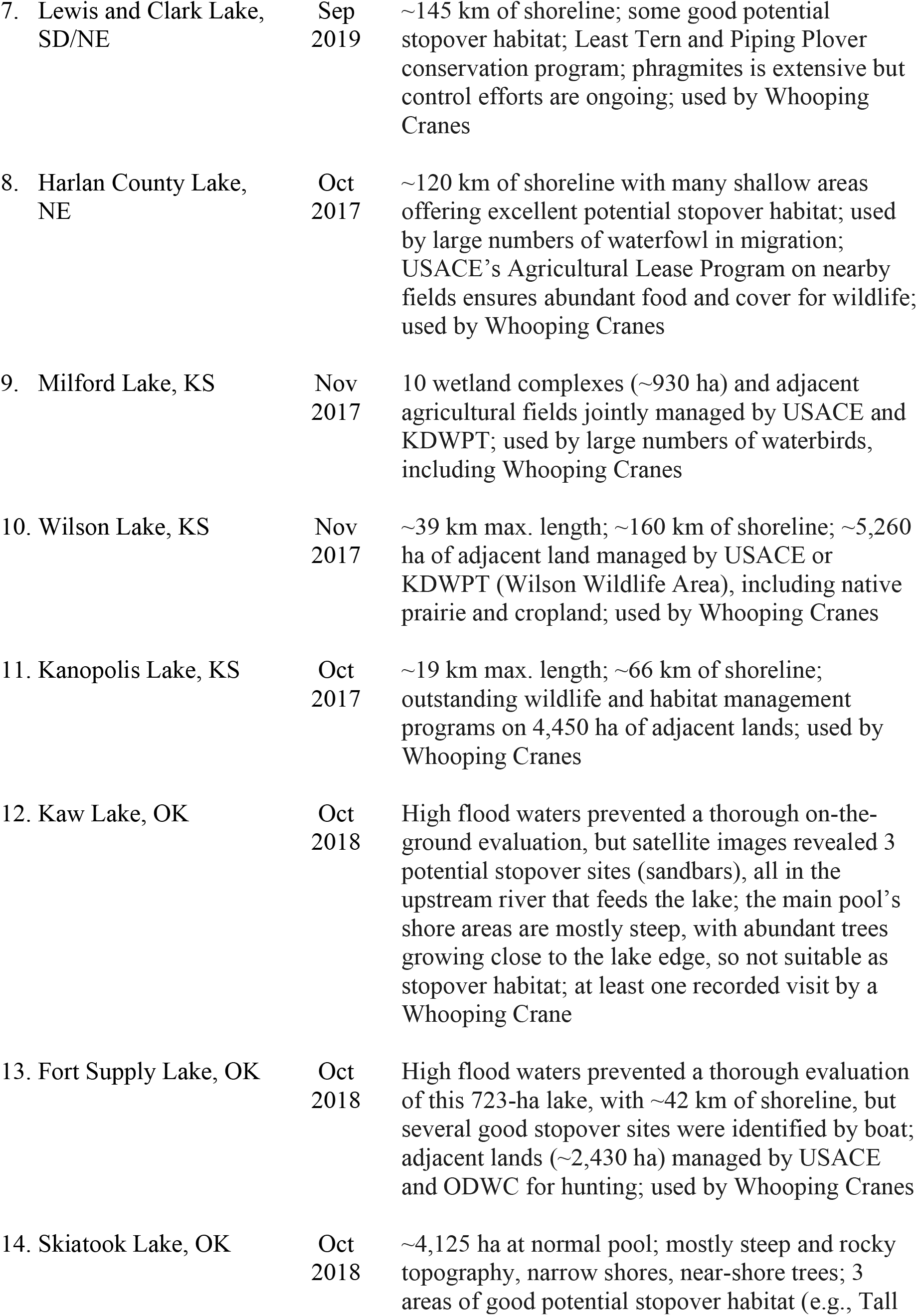
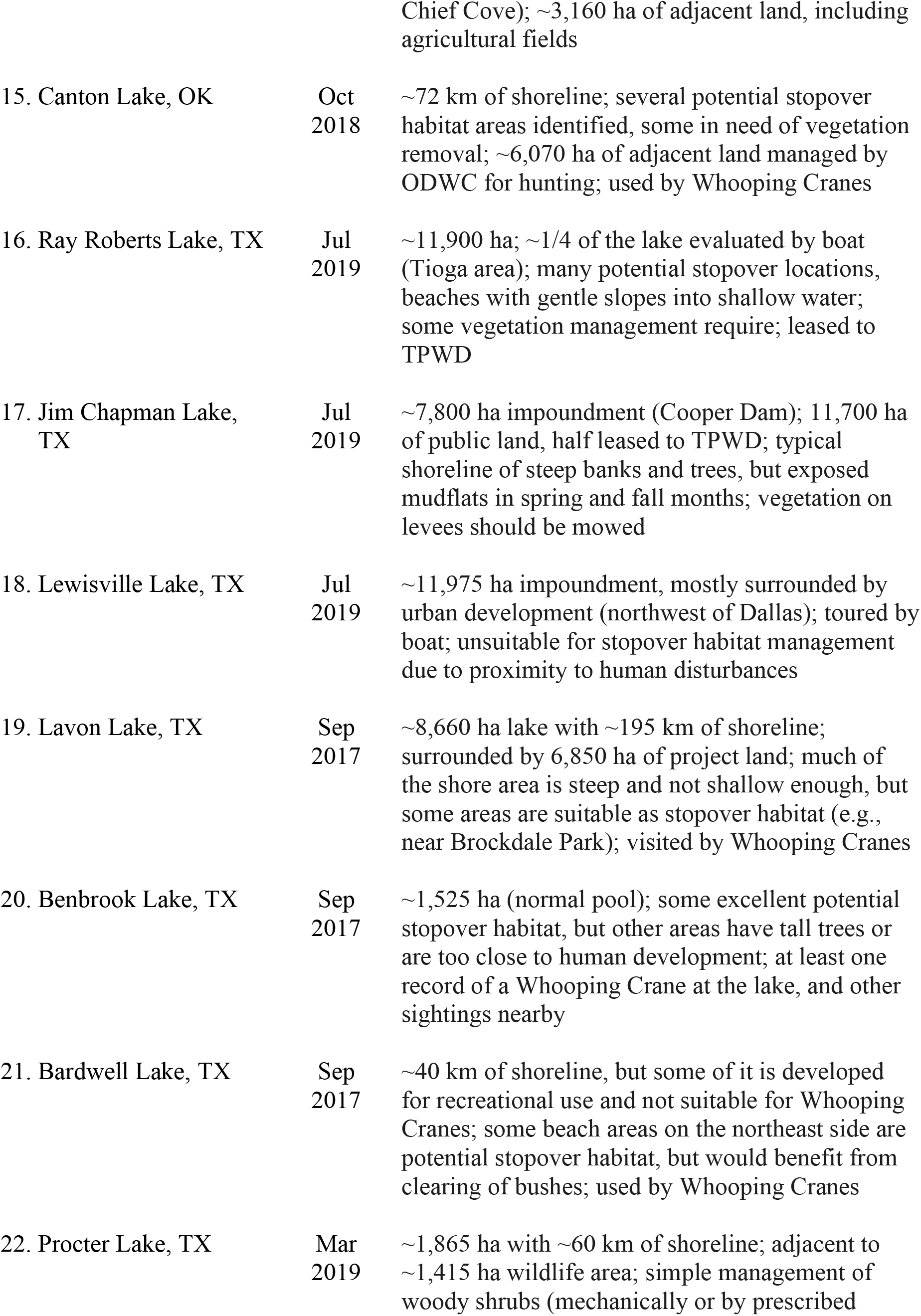
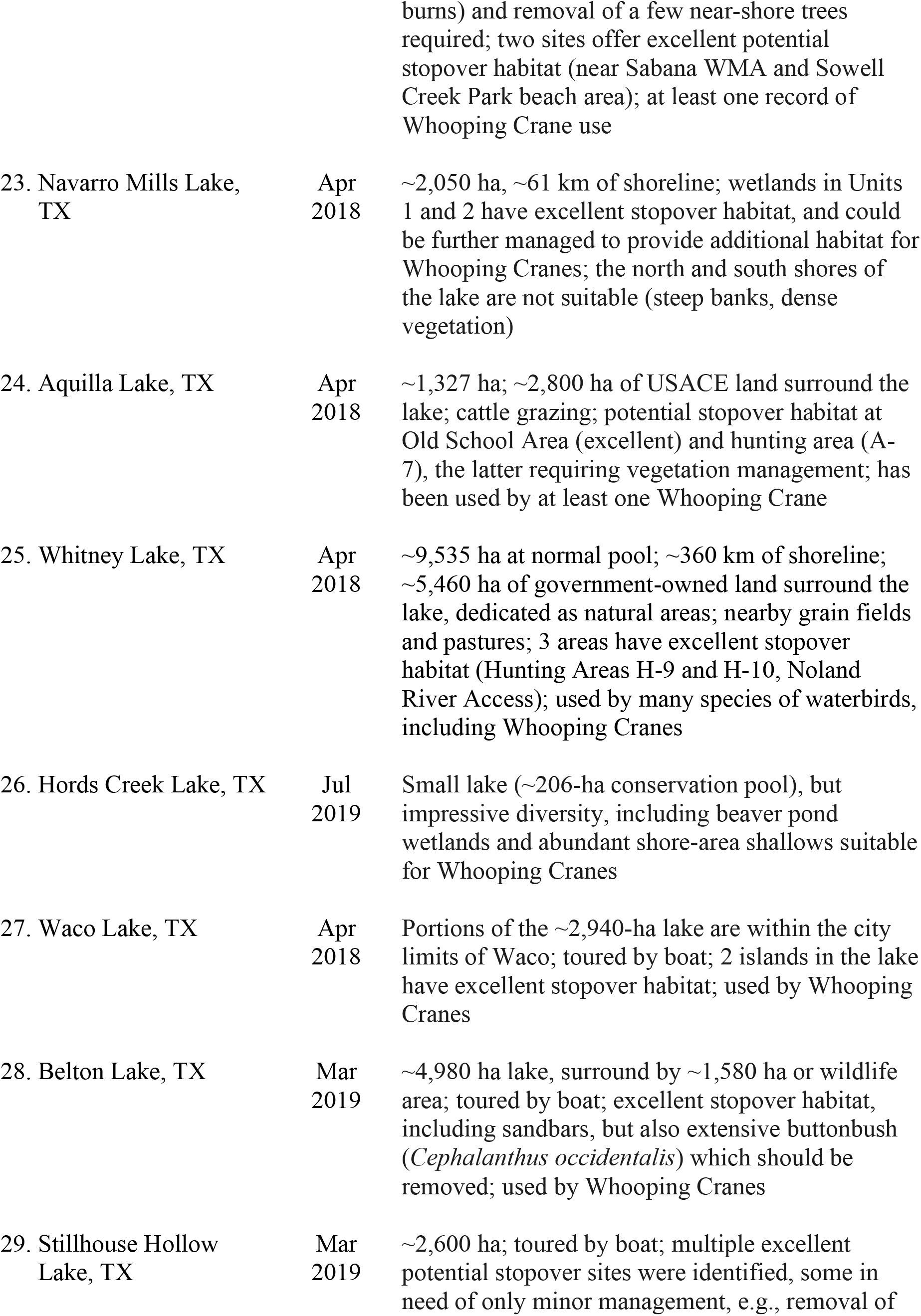
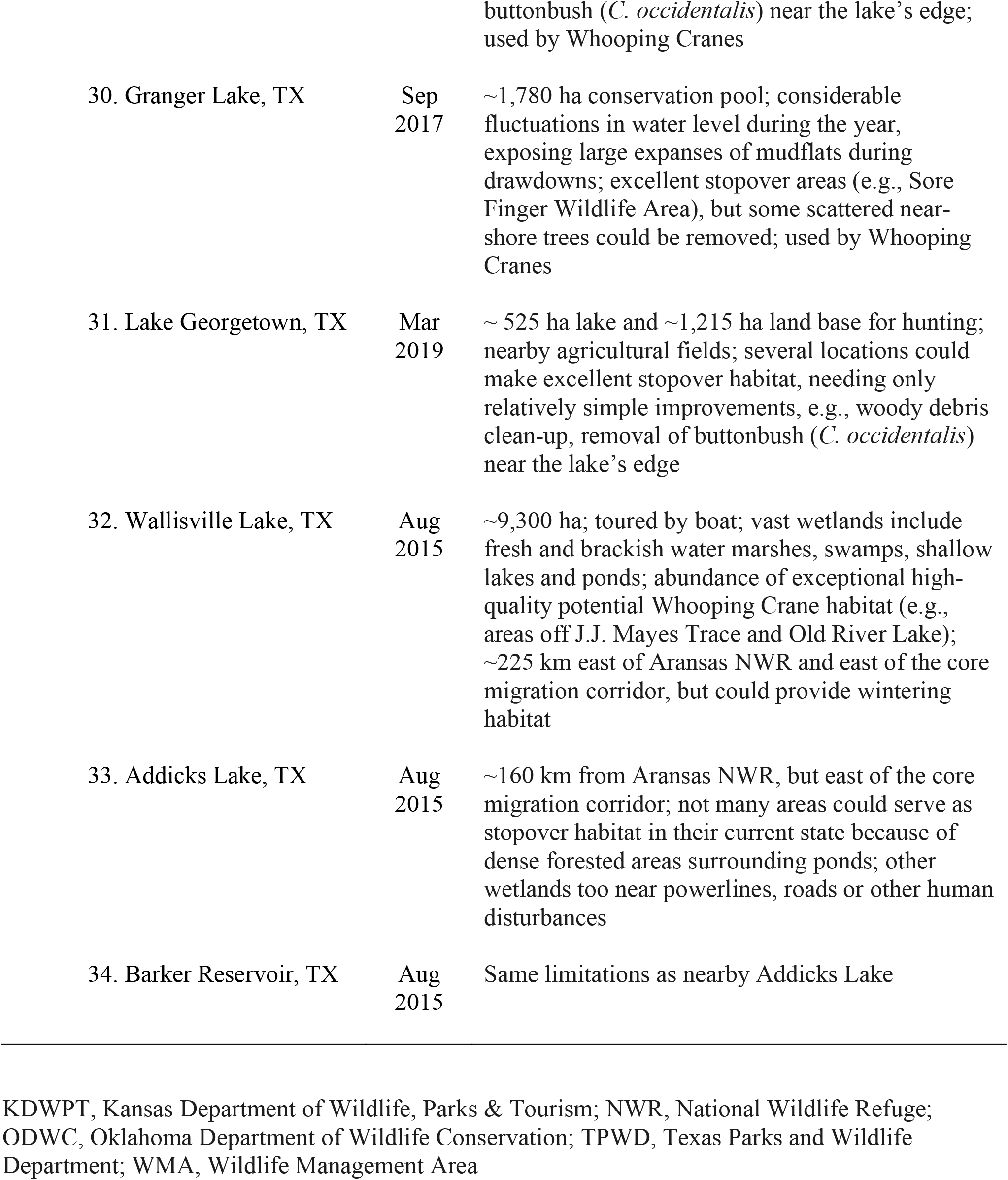
USACE lakes visited by FOTWW to identify potential stopover locations that could be managed for migrating Whooping Cranes of the Aransas-Wood Buffalo Population. Lakes are listed by state and from highest to lowest latitude.

## RESULTS

FOTWW conducted field trips on 34 USACE properties in seven states from August 2015 to September 2019 **(see Table 1; Figures 2-4)**. Three USACE lake properties (Addicks, Barker and Wallisville, all near Houston, Texas; Figure 4) which were included as ‘military bases’ in McConnell (2018) are mentioned again here for completeness.

**Figure 2.**
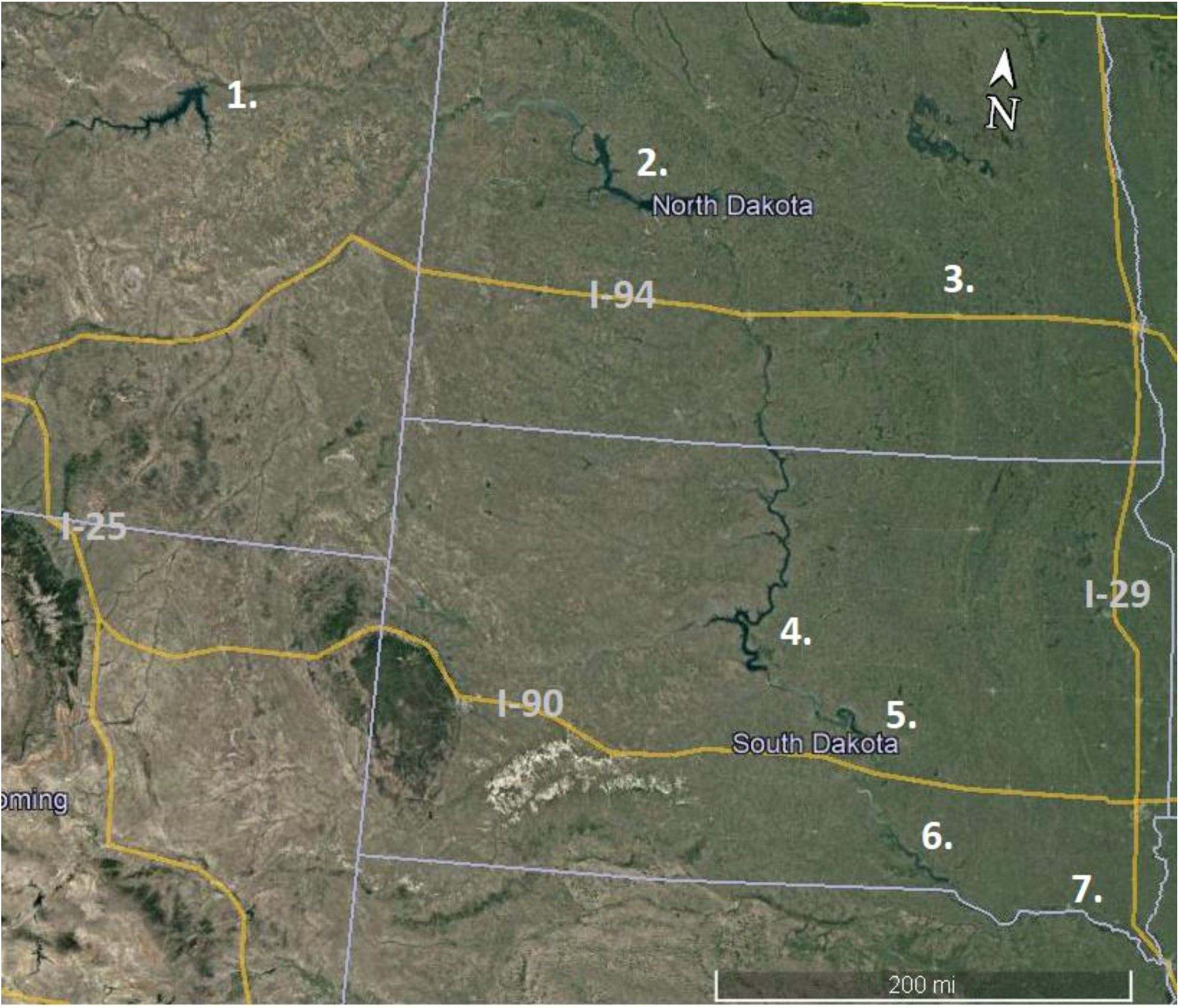
Field visit sites 1-7 in Montana, North Dakota and South Dakota. The numbers on the map correspond to the numbered USACE lake locations in Table 1. Interstate highways are labeled. Mapping source: 2020 Google, Image Landsat Copernicus. Scale bar = 320 km.

**Figure 3.**
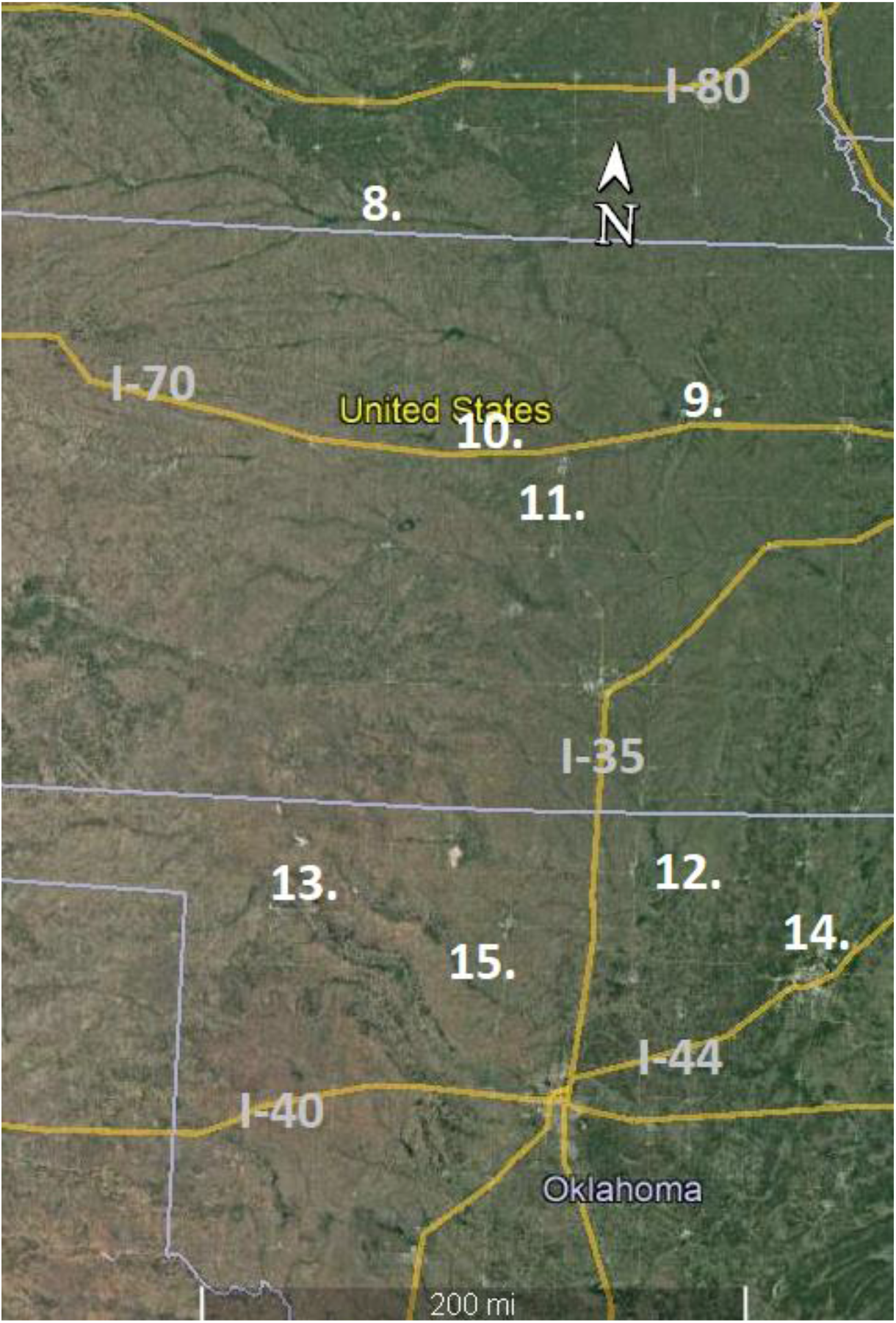
Field visit sites 8-15 in Nebraska, Kansas and Oklahoma. The numbers on the map correspond to the numbered USACE lake locations in Table 1. Interstate highways are labeled. Mapping source: 2020 INEGI, 2020 Google, Image Landsat Copernicus. Scale bar = 320 km.

**Figure 4.**
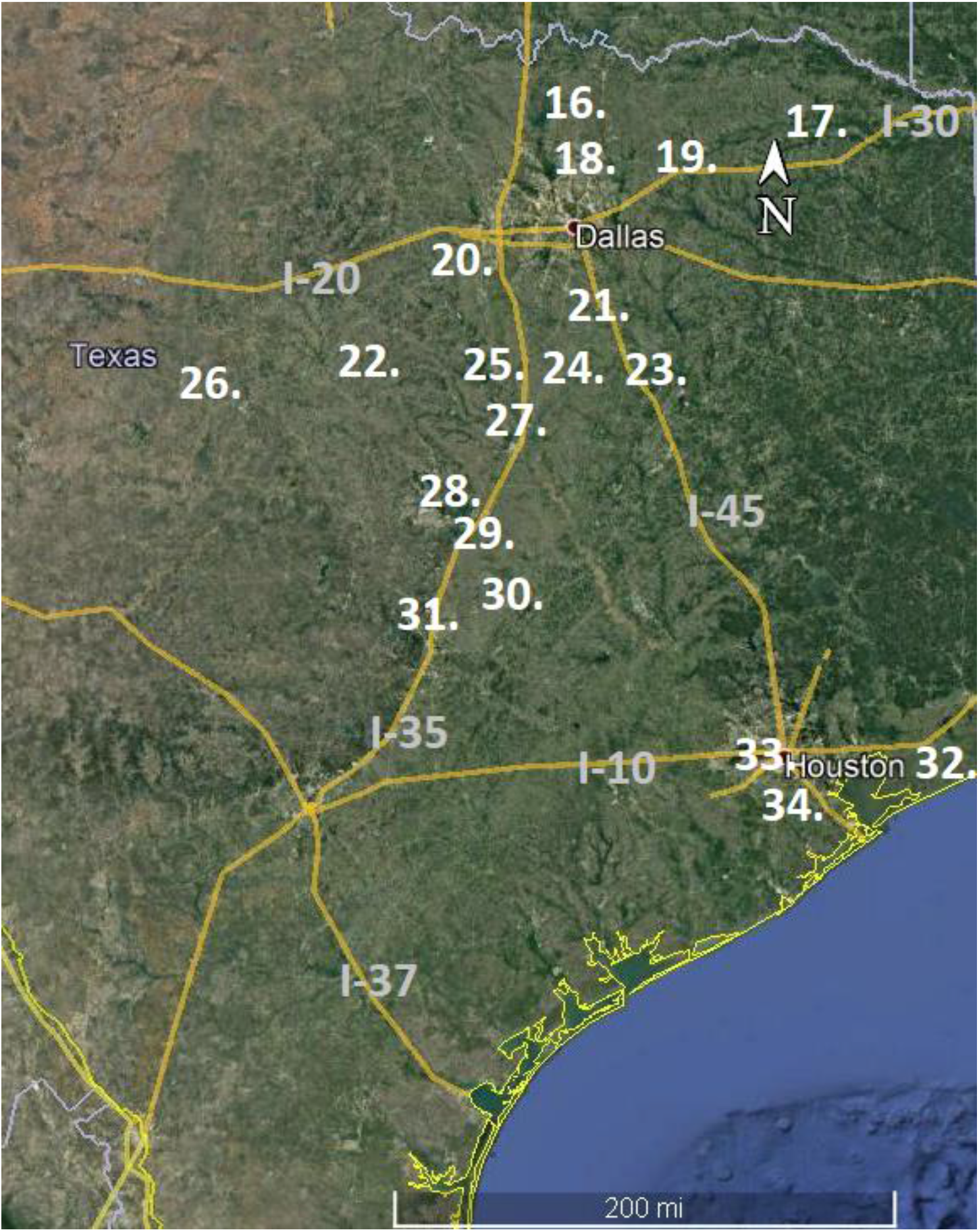
Field visit sites 16-34 in Texas. The numbers on the map correspond to the numbered USACE lake locations in Table 1. Interstate highways are labeled. Mapping source: 2020 INEGI, 2020 Google, Image Landsat Copernicus, Data SIO-NOAA, U.S. Navy, NGA, GEBCO. Scale bar = 320 km.

FOTWW discussed Whooping Crane biology, habitat management needs and specific management practices with USACE (and sometimes state) wildlife biologists during the field trips. We then developed detailed management recommendations for each lake to protect, improve or develop potential Whooping Crane stopover habitats and provided detailed reports for each USACE property explaining our management recommendations (summarized in Table 1). Copies of FOTWW recommendations, in the form of written reports, were provided to all personnel involved.

Of the 34 lakes we visited, many had sites that already met FOTWW stopover habitat criteria or needed only inexpensive management practices to become suitable for migrating Whooping Cranes, e.g., by cutting dense vegetation around the edge of the lakes (e.g., Canton Lake, OK, Procter Lake, TX, Belton Lake, TX, Stillhouse Hollow Lake, TX, among others; Table 1). Importantly, FOTWW estimated that 624 potential stopover wetland habitats on these 34 lakes could be used by Whooping Cranes by undertaking varying degrees of habitat management. Indeed, we learned retrospectively that many of the lake properties we visited have records of Whooping Crane use (Table 1), thereby supporting the efficacy of our approach. However, some lakeside locations are not useful for Whooping Cranes because of proximity to human disturbance (e.g., Lewisville Lake, TX); or steep and rocky shorelines (e.g., Skiatook Lake, OK); or cattails, bushes (e.g., buttonbush, *Cephalanthus occidentalism* and trees are currently thick along the shore areas (see Table 1). On these latter locations, FOTWW recommends that they be managed for other wildlife species that prefer dense vegetative cover. Indeed, FOTWW contends that it is not necessary or desirable to modify or manage all wetlands for Whooping Cranes, but rather to focus on a subset with the best habitats and surrounding landscape characteristics.

## DISCUSSION

The development and management of stopover habitat for AWBP Whooping Cranes as recommended by FOTWW would not be expensive, because USACE already owns the land and waters where these stopover habitats are located. As an important outcome of our site visits, USACE officials were encouraged to protect and manage the identified wetlands as part of the USACE Environmental Stewardship Program. All USACE personnel advised that they intended to implement our recommendations over time as funding and time permits. Indeed, the USACE has environmental laws and regulations that it must follow (McConnell, 2018). For example, in accordance with the Endangered Species Act of 1973, as amended, the Army must assist in recovery of all listed threatened and endangered species and their habitats under the Army’s land management authority. Importantly, the Sikes Improvement Act of 1977 (16 U.S.C.670) requires the Secretary of Defense to carry out a program to provide for the conservation and rehabilitation of natural resources on lands used for military mission activities. Furthermore, the Migratory Bird Treaty Act (16 U.S.C.703-712) requires protection of migratory birds. Based on FOTWW observations, the USACE personnel we met with are using all these legal authorities to manage lands in a manner beneficial to many species of wildlife, including Whooping Cranes.

Since we completed the USACE phase of our evaluation, about one quarter of the land managers have contacted FOTWW to discuss management practices in more depth. Moreover, personnel at the USACE Engineer Research and Development Center’s Environmental Laboratory and USACE Headquarters have begun working closely with the US Geological Survey to analyze multiple years and thousands of GPS satellite tag locations to confirm significant use of USACE land and water as stopover habitat within the AWBP Whooping Crane migration corridor. In support of the MOU and in accordance with USACE responsibilities, the USACE has committed to identifying measures to maintain existing stopover habitat, improving habitat where possible, coordinating with the USFWS under the Endangered Species Act in the context of potential habitat improvement projects, and annual monitoring of habitat use by Whooping Cranes to evaluate the effectiveness of habitat maintenance and restoration projects. Moreover, because the lands and waters are USACE properties, the cost of stopover habitat enhancement and management will be relatively minor.

So, what did FOTWW accomplish on the USACE lake properties? As with the military bases and Indian Reservations (McConnell, 2018), awareness and interest in Whooping Cranes by natural resource personnel was significantly increased, as was their desire to help endangered Whooping Cranes. USACE personnel were encouraged to protect and manage several hundred potential stopover wetlands identified by FOTWW, thus targeting some of the major unmet objectives described in the Whooping Crane Recovery Plan, which include identifying, protecting, managing, and creating stopover habitat for Whooping Cranes. FOTWW contends that wild AWBP Whooping Cranes are capable of taking care of themselves, with two exceptions. They need people to protect their wetland habitats and to protect them from gunshot.

## ACKNOWLEDGMENTS

I sincerely appreciate the interest and cooperation of all the outstanding USACE personnel during our evaluation of potential AWBP Whooping Crane stopover habitats on USACE properties. I am particularly grateful to David Hoover, Conservation Biologist, Kansas City, Missouri, USACE, who arranged our visits and provided documents that assisted in our evaluations. David also graciously prepared the maps used in figures 2, 3 and 4. Lastly, I thank Daryl S. Henderson, Ph.D., for editing assistance on this paper. The findings and conclusions in this paper are those of FOTWW.

## Notes

### Competing Interest Statement

The authors have declared no competing interest.

